# The non-redundant nature of the Axin2 regulatory network in the canonical Wnt signaling pathway

**DOI:** 10.1101/2021.04.21.440851

**Authors:** Ana R Moshkovsky, Marc W Kirschner

## Abstract

Axin is one of two essential scaffolds in the canonical Wnt pathway that converts signals at the plasma membrane to signals inhibiting the degradation of β-catenin, leading to its accumulation and specific gene activation. In vertebrates there are two forms of Axin, Axin1 and Axin2, which are similar at the protein level and genetically redundant. We show here that differential regulation of the two genes on the transcriptional and proteostatic level confers robustness and differential responsiveness that can be used in tissue specific regulation. Such subtle features may distinguish other redundant gene pairs that are commonly found in vertebrates through gene knockout experiments.

**Significance Statement:** The mystery of two functionally redundant Axin genes in all vertebrates are can now be explained by the demonstration that they form a nested proteostatic and transcriptional feedback system that confers regulatory options in different developmental settings, a form of dynamic versatility that may explain the widespread occurrence of closely related seemingly redundant genes with similar functions.

## INTRODUCTION

The Wnt signaling pathway is present in all metazoan animals where it plays an important role in global embryonic axis formation as well as in local patterning and differentiation in many tissues (1, 2). It is also of importance in human disease, such as colorectal cancer and hepatocellular carcinoma. The pathway involves multiple inputs from secreted Wnt protein ligands (17 different ones in humans) through Frizzled receptors (10 in humans) to various outputs. The so called canonical pathway is best known in embryology and cancer for its control of the level of β-catenin, which activates transcriptional targets. The rise in β-catenin is not caused by activating transcription or translation, but by inhibiting the high steady state level of β-catenin degradation, which is regulated by β-catenin phosphorylation (3). The pathway by which the Wnt ligand inhibits phosphorylation of β-catenin itself is complex and still not completely understood.

Central to the canonical Wnt pathway are two scaffold proteins, Adenomatous polyposis coli (APC) and Axin, both essential. The scaffolds promote phosphorylation of β-catenin, and cause its degradation. The Axin scaffold is especially important by co-localizing kinases and β-catenin (4). More than 20 years ago it was discovered that there are two forms of Axin (Axin1 and Axin2) in vertebrates, whereas insects and nematodes have only one. In cultured cells Axin2 can fully substitute for Axin1 but its use as a backup can scarcely be its biological function or else every essential gene should have a second copy. For this reason Axin2 is believed to possess some specific biochemical property, such that it regulates the duration or intensity of the Wnt signal (5-7). Intensity and duration of signaling should be reflected in different kinetic properties of the pathway. Hence, we wish to develop a clearer understanding of the kinetics of Axin2 and its effects on the dynamics of β-catenin. Using quantitative kinetic assays and gene knockdowns we show that Axin2 conveys previously unappreciated attributes to the Wnt response, endowing it with new forms of regulation, thereby explaining why Axin2 is dispensable in some circumstances and required in others.

## RESULTS

### The response of β-catenin to Wnt stimulation is insensitive to Wnt-induced Axin2 expression

We examined the effect of Axin2 silencing in the response of β-catenin to Wnt stimulation. We had previously analyzed the dynamics of β-catenin in Wnt3A-stimulated RKO cells by steady state and pseudo steady state kinetics (3). We find no effect of Axin2 silencing on the dynamics of total or on the so called “active” non phospho form of β - catenin (fig. 1A). Although the special biological significance of unphosphorylated β-catenin is questionable, increase of this non-phosphorylated species, rather than total β-catenin, had been previously reported in Axin2 knock-out mice (7).

**Fig. 1.**
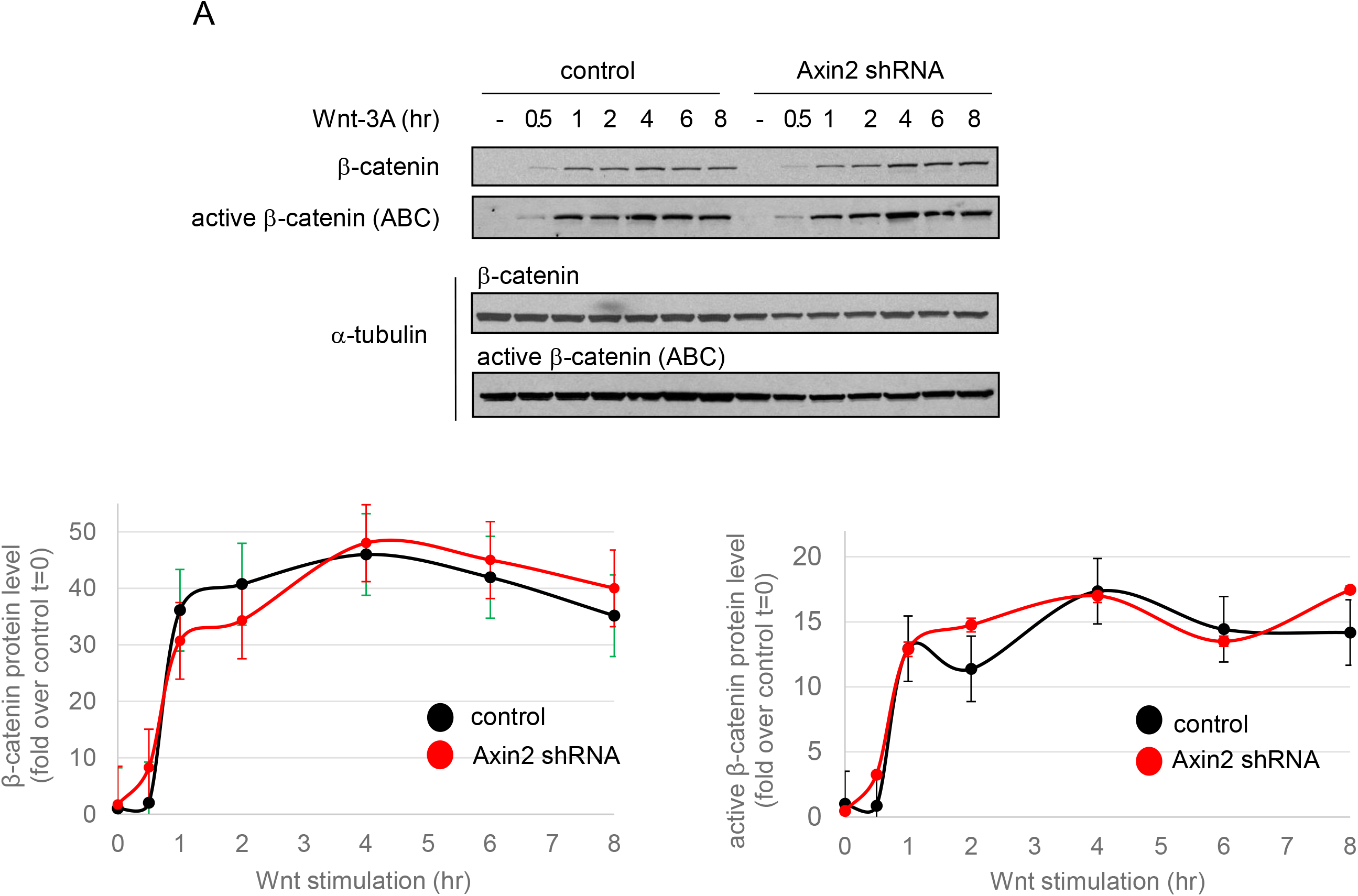
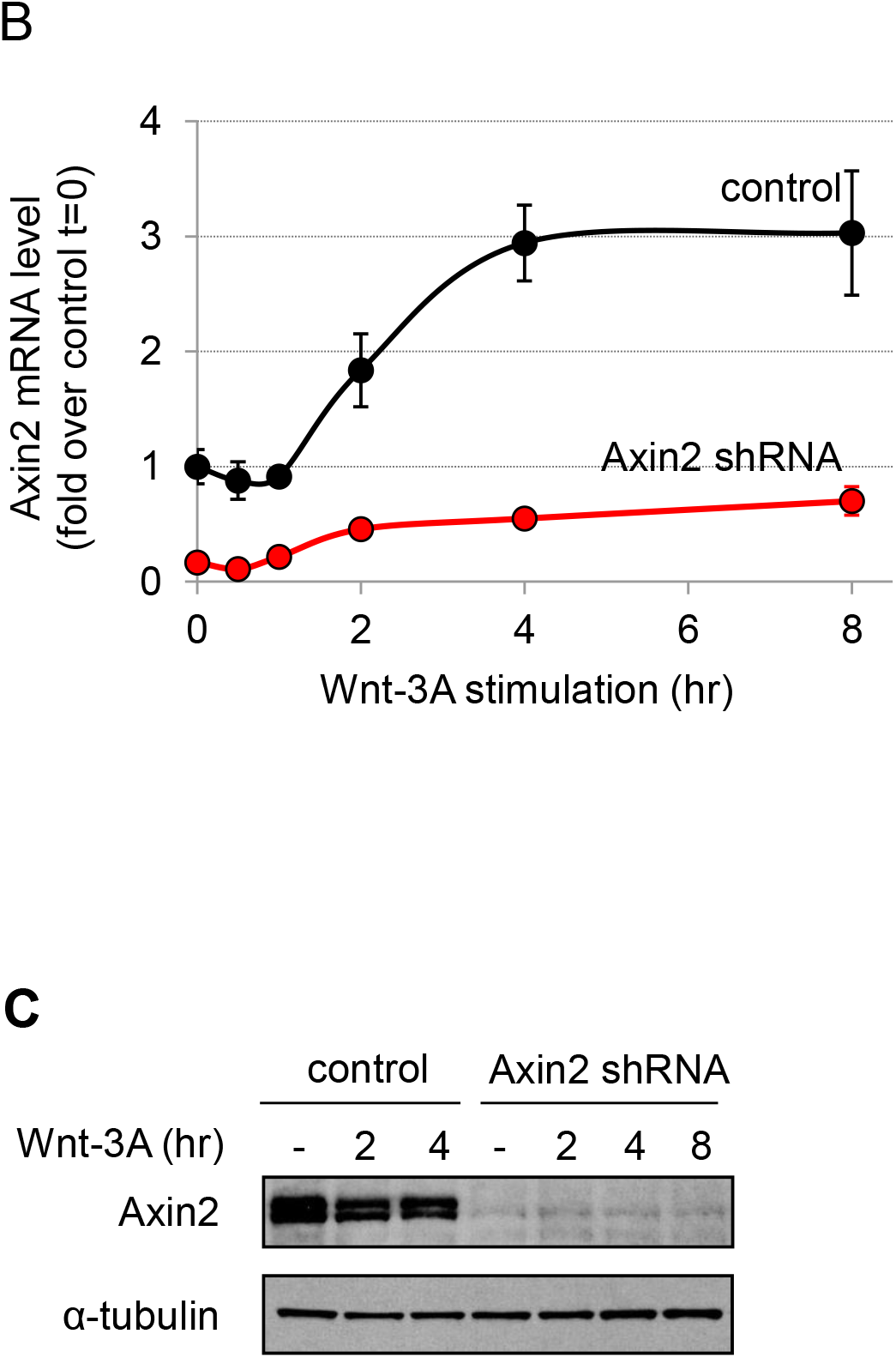
The RKO cell response to Wnt-3A stimulation remains unaffected after Axin2 silencing. (**A**) Western blotting and protein quantification (N=3) of β-catenin, and active β-catenin proteins in RKO control cells and RKO cells expressing Axin2 shRNA, after Wnt-3A stimulation for 0 to 8 hours. (**B**) RT-PCR (N=3) of Axin2 in RKO control cells and RKO cells expressing Axin2 shRNA. (**C**) Western blotting of Axin2 in RKO control and Axin2-shRNA expressing cells.

The lack of any measurable effect of Axin2 silencing on β-catenin accumulation in Wnt-stimulated cells was not due to ineffective silencing Axin2 expression. Axin2 mRNA levels over the course of eight hours of the Wnt response (fig 1B), were reduced by 80-85% in Axin2 shRNA-treated RKO cells; Axin2 protein levels were reduced more than 95% (fig. 1C), both before and after Wnt-3A stimulation. Therefore, this lack of effect cannot be explained by failure to silence Axin2 expression. Though these results suggest that Axin2 plays no essential role in the dynamics of β-catenin in the Wnt response under conditions where Axin1 is present, they do not prove that Axin2 is not an essential gene under some conditions, although there is no known situation where only Axin2 is present. Despite this seeming lack of effect in cultured cells, a deletion of the Axin2 gene has phenotypic effects in animals, such as significant overgrowth of bone (7) and tooth agenesis (8), suggesting either that we had not probed deeply enough or possibly that Axin2 has an important role outside the Wnt pathway.

### Wnt3A stimulation activates Axin2 transcription and simultaneously destabilizes Axin2 protein

To pursue further a role for Axin2 in the Wnt pathway we studied how Axin 2 responds to a Wnt signal. As previously found (5), Wnt3A stimulation activated Axin2 mRNA transcription, elevating the levels 2 fold in HEK293T cells, 3 fold in RKO and 13 fold in U-2 OS cells (fig. 2A). This rise in Axin2 mRNA most likely is a response to β-catenin, since the accumulation of Axin2 occurs with similar kinetics to the increase in the level β-catenin (fig. 2B).

**Fig. 2.**
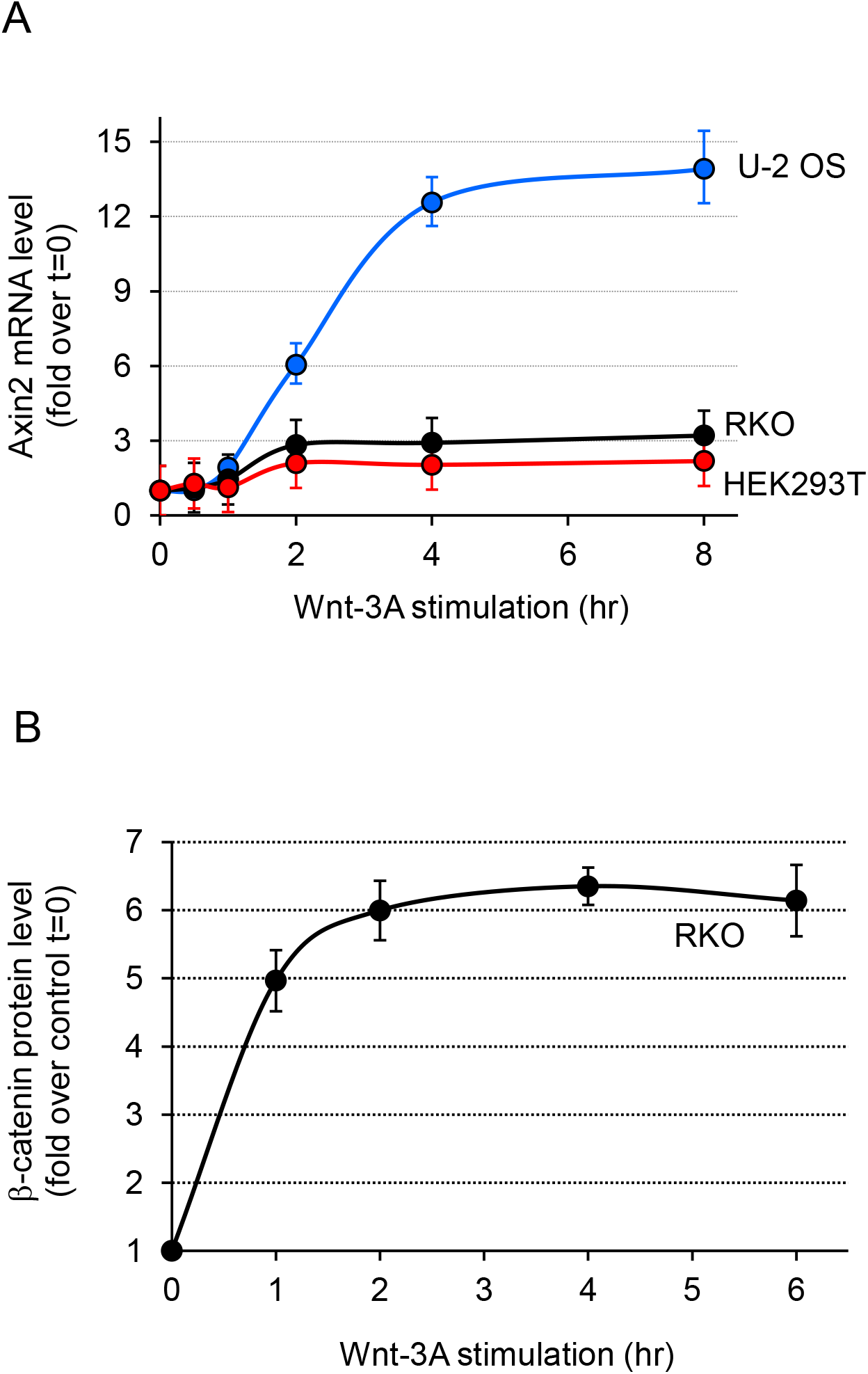
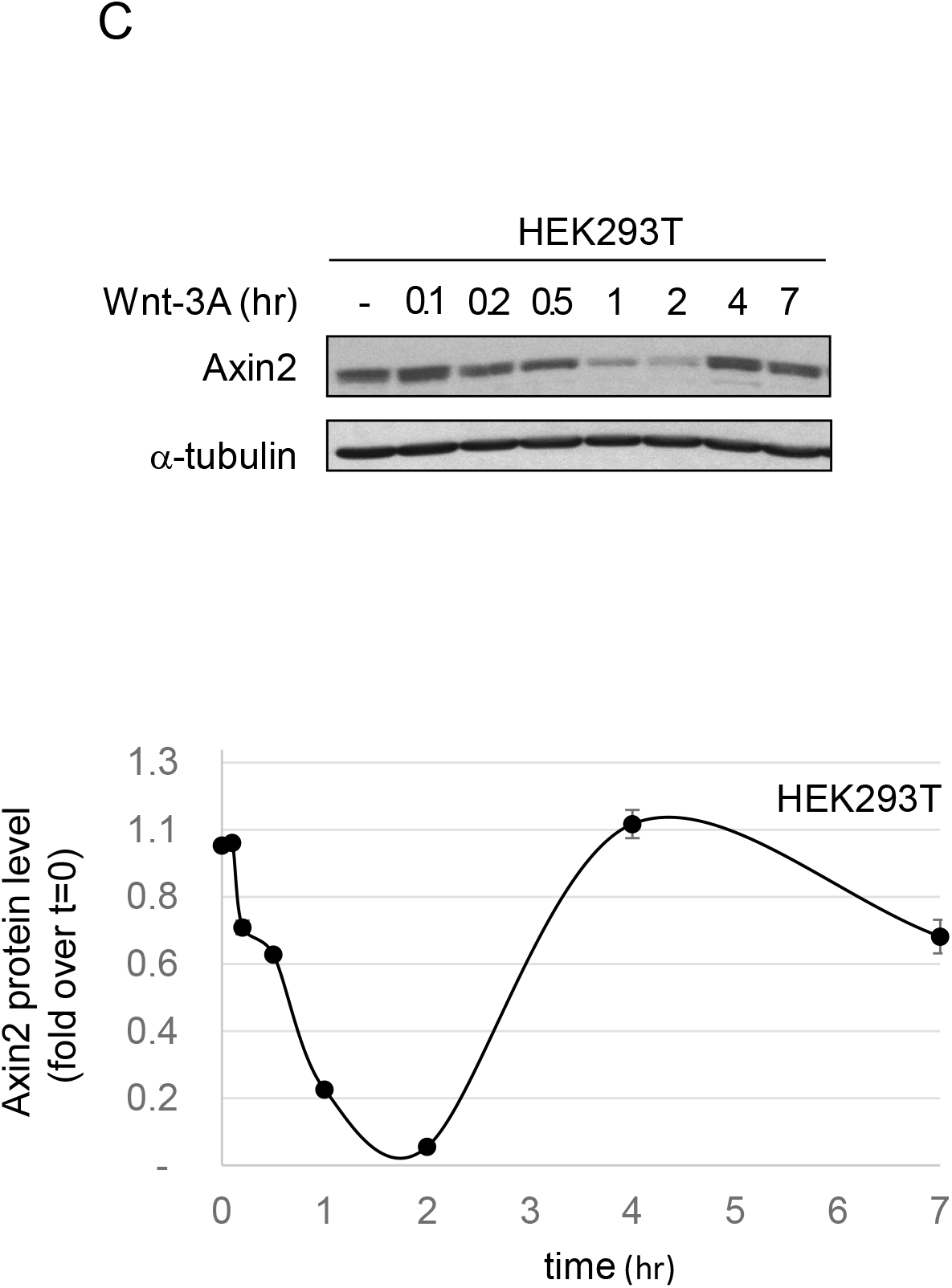
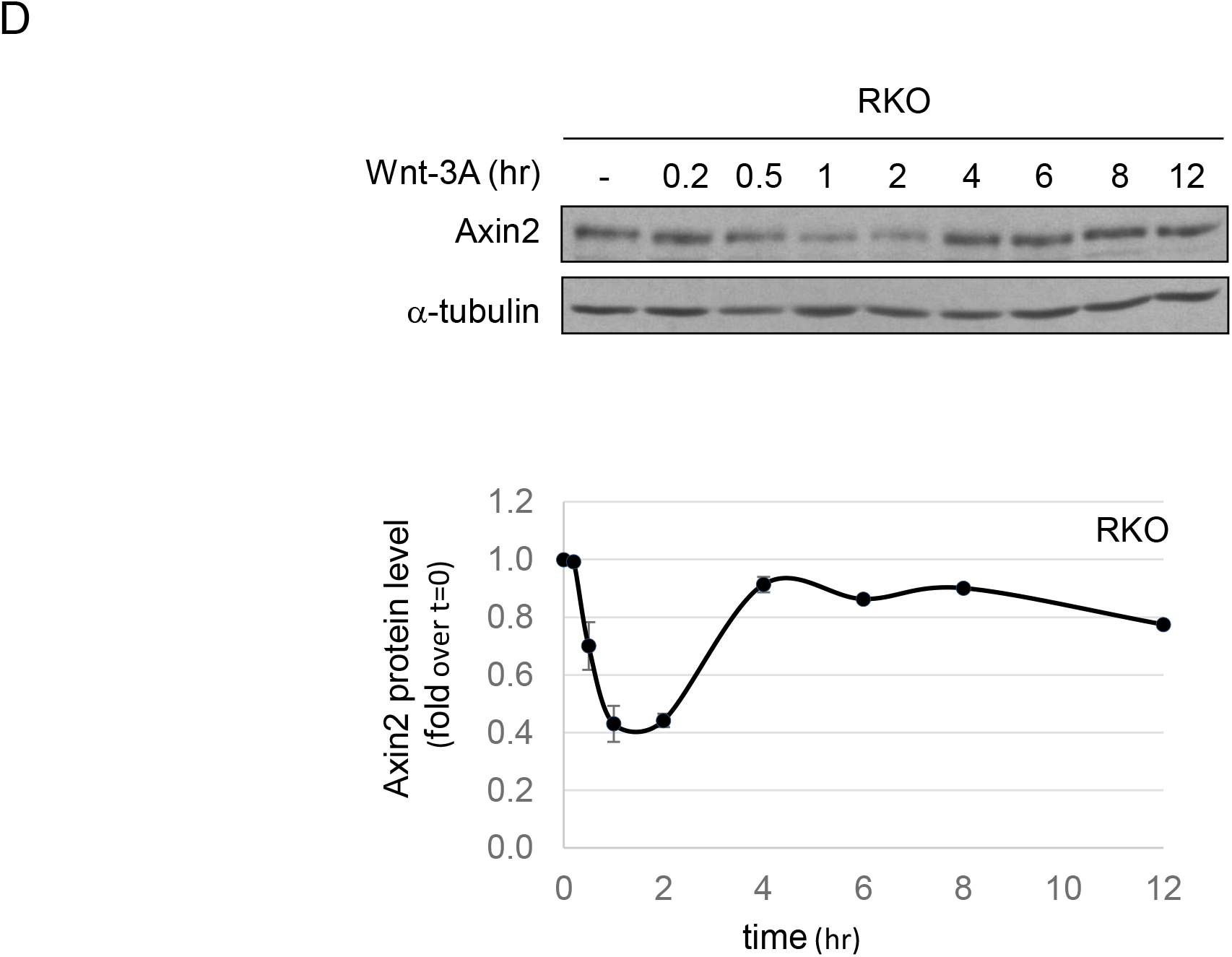
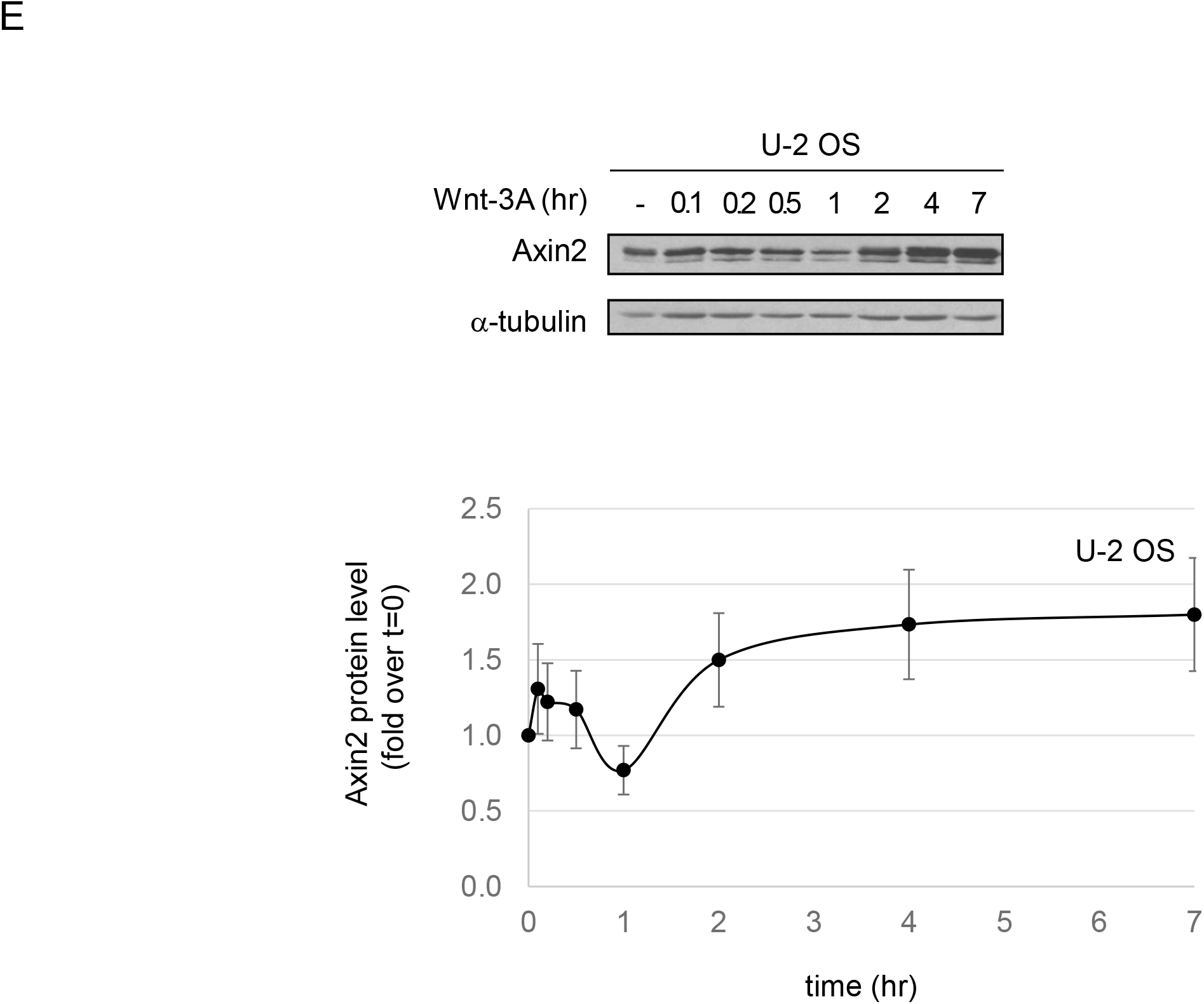
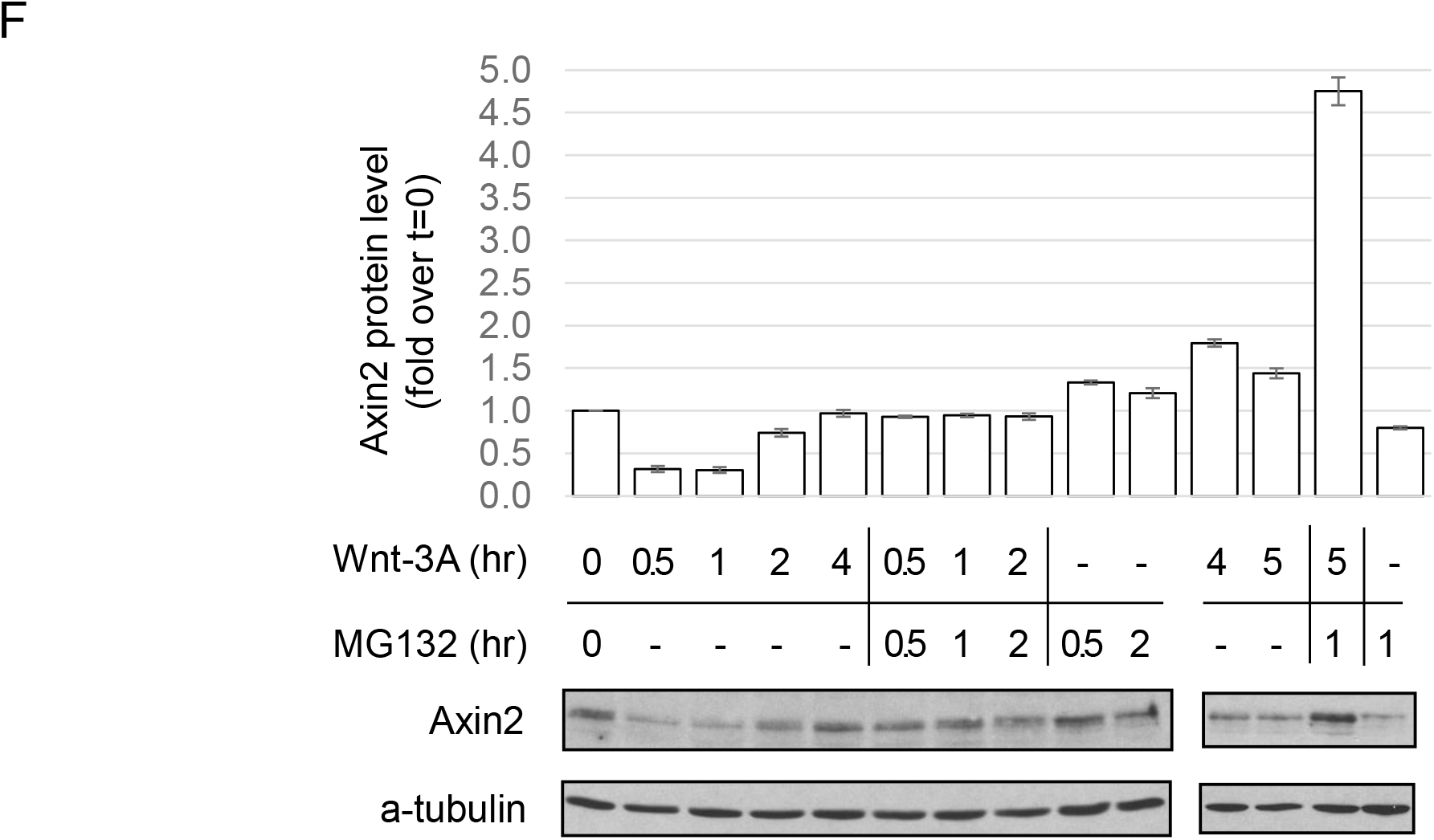
Axin2 protein is degraded in response to Wnt-3A stimulation. (**A**) RT-PCR (N=3) of Axin2 in RKO, HEK293T and U-2 OS cells. (**B**) Protein quantification of β-catenin in RKO control and Axin2 shRNA expressing cells. (**C-E**) Western blotting and protein quantification (N=3) of Axin2 in HEK293T, RKO and U-2 OS cells after Wnt-3A stimulation. (**F**) Western blotting and protein quantification (N=3) of Axin2 in Wnt3A-stimulated RKO cells with or without treatment with the proteasome inhibitor MG132.

However, the protein level of Axin2 does not parallel the monotonic increase in mRNA transcript levels, as shown in Figure 2C. Rather, there is an initial steep 2 to 4-fold decrease of Axin2 protein in different cell lines in the first hour of Wnt exposure, followed by a recovery in Axin2 to the initial protein level in HEK293T and RKO cells (fig. 2C, 2D and 2E); in U-2 OS cells there is 2-fold overshoot. These dynamics imply that in response to Wnt, there is a marked increase in Axin2 protein degradation, which is ultimately overwhelmed by the transcriptionally driven increase in Axin2 protein. To test that this initial drop of Axin2 is due to protein degradation, we treated cells with the proteasome inhibitor MG132 (fig. 2F). MG132 completely prevented the loss of Axin2 protein, clearly showing that Axin2 is destabilized under Wnt stimulation. Furthermore, the turnover of Axin2 during the first 2 hrs. seems to be very fast, as MG132 treatment increased the level of Axin2 protein within one hour. Destabilization of Axin2 continues at least for 5 hours after addition of Wnt (fig. 2F). In RKO and HEK293T cells, increased Axin2 transcription drives Axin2 protein synthesis and restores the initial loss of Axin2 protein. However, in U-2 OS cells, the increase in Axin2 transcription was 5 times higher than in RKO and HEK293T (fig. 2A) causing a higher accumulation of Axin2 protein.

To test whether Axin2 has any role in regulating β-catenin levels in the absence of Wnt we asked how loss of Axin1 would affect Axin 2. We would expect that Axin1 specific shRNA treatment would stabilize β-catenin and the rise in β-catenin would activate Axin2 transcription. As predicted, when we silence Axin1 expression, β-catenin protein levels increased 4.5 fold (fig. 3A) and Axin2 protein levels increased 5.3 fold (fig. 3B). To understand how these levels change in response to both Axin knockdowns in the absence of Wnt, we silenced both Axins; β-catenin climbed to a level 9.8 times the original level (fig. 3A), compared to 5.3 fold for the suppression of Axin1 alone, demonstrating that Axin2 makes a very significant contribution to negatively regulating β-catenin protein levels in the absence of Axin1. Despite that contribution, the knockdown of only Axin2 did not change the levels of β-catenin. The effects of Axin1 and Axin2 knockdowns are clearly not additive. To summarize, the Axin2 knockdown had no measurable effect, while the Axin1 knockdown raised the level of β-catenin 4.5 fold and both raised the level 9.8 fold, suggesting that Axin1 and Axin2 must interact. We further studied the additivity using RKO cells expressing a firefly luciferase β-catenin transcriptional reporter (9). Fig. 3C shows that although Axin1 silencing increased the luciferase activity and Axin2 silencing had a similar effect, the luciferase bioluminescence was still 200 times lower to the case when both Axin1 and Axin2 were both silenced. This would not be the case if Axin2 made a negligible contribution or even if the contributions were equal. It can only be explained by some cross suppression.

**Fig. 3.**
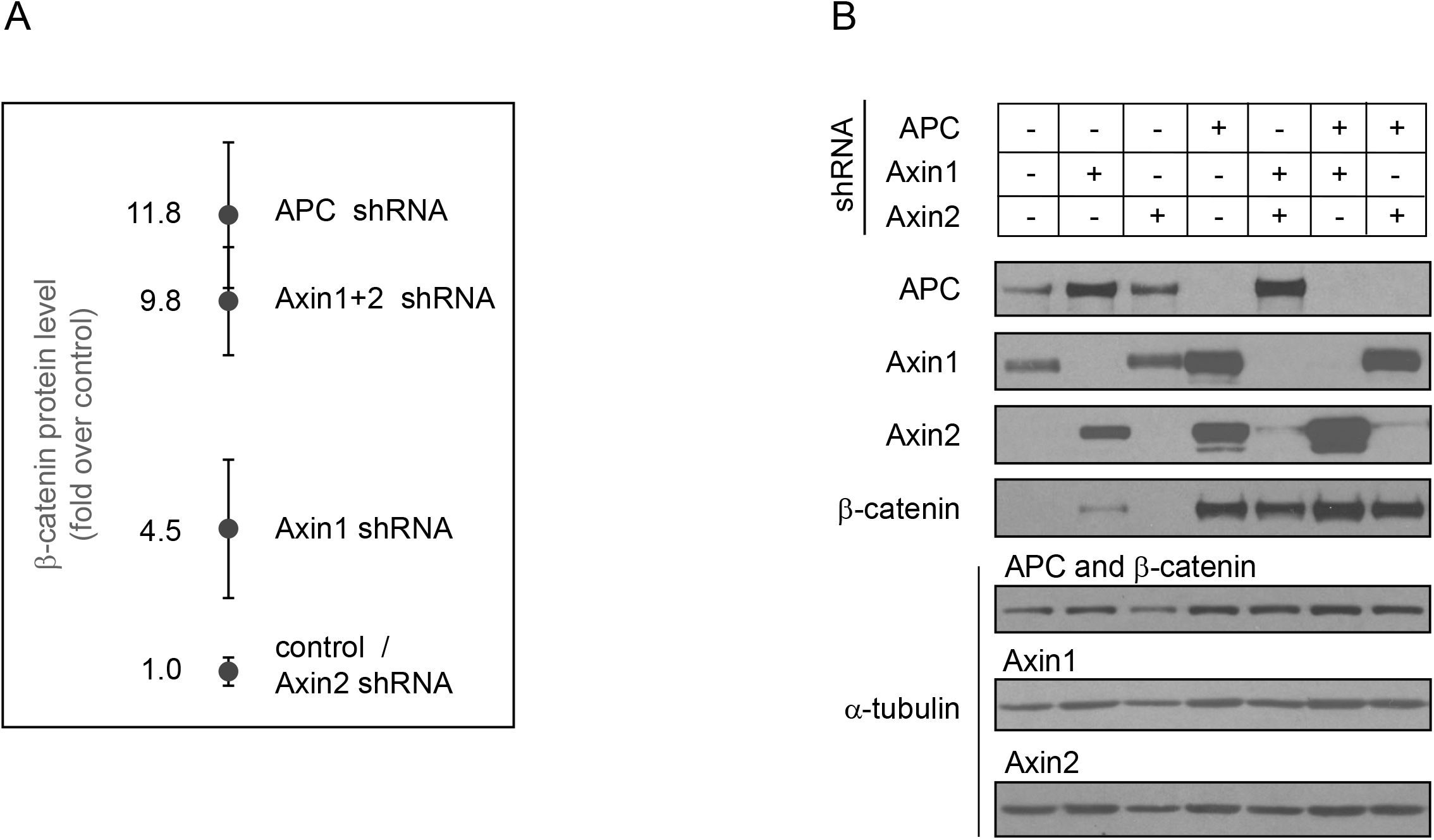
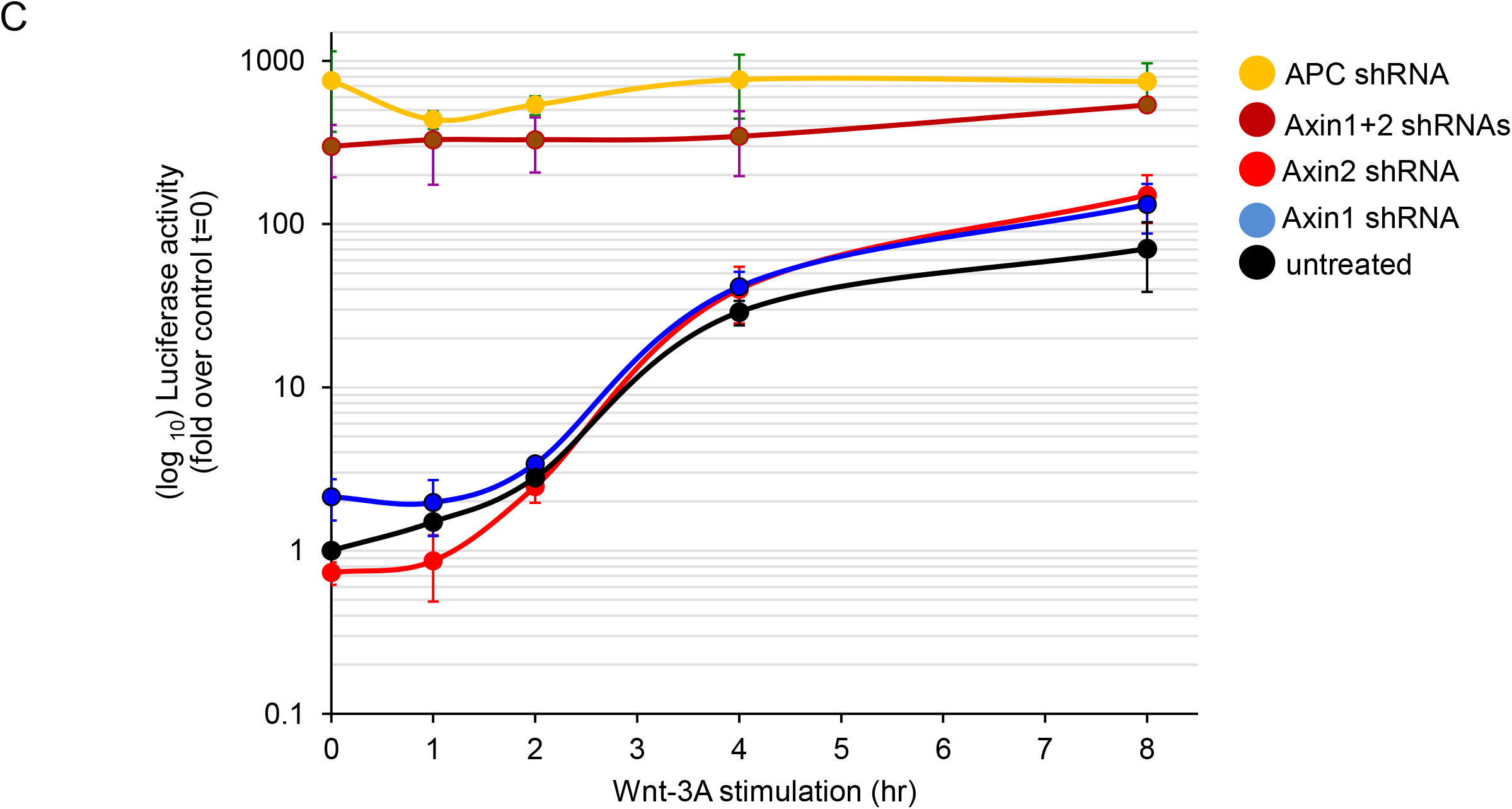
Axin2 protein expression partially compensates loss of Axin1. (**A**) Protein quantification (N=3) of β-catenin in RKO control cells after Axin2, Axin1, Axin1+Axin2 and APC knock down. (**B**) Western blotting of APC, Axin1, Axin2 and β-catenin under the indicated conditions of shRNA silencing. α-tubulin is shown as a loading control. (**C**) Firefly luciferase activity (N=3) in RKO cells stably expressing a β-catenin-activated reporter gene under the indicated knock down conditions after Wnt-3A stimulation.

## DISCUSSION

Axin2 plays a distinct but subtle role in regulating the level of β-catenin in the presence and absence of Wnt. However, in a Wnt-stimulated cell the effective suppression of Axin2 does not measurably change the rate or extent of β-catenin accumulation in response to Wnt3A. As redundant as Axin2 seems from knock down experiments, it is nevertheless hard to ignore that Axin2 is one of the most responsive downstream transcriptional targets of β-catenin, even though other core genes in the Wnt pathway, such as APC or Disheveled are not regulated in this manner. The well reported Axin2 transcriptional response was a clue to us that despite the negligible effect of the knockdown of Axin2 in cells, we may be missing an important regulatory role for Axin2 in the Wnt response. The dynamics of Axin2 mRNA are dramatic. It accumulates 2 to 12 fold in different cell lines over 4 hrs. with a very small delay (approximately 1 hr.) after Wnt stimulation. From the rapid mRNA accumulation, we might have expected a rapid down regulation of β-catenin and perhaps only a transient peak of Wnt regulated gene expression, but this is not what is seen. Despite the precipitous rise in Axin2 mRNA after Wnt stimulation in RKO and HEK293 cells, there is no increase in Axin2 protein levels until 4 hours; (there is only a 2-fold increase in Axin2 protein in U-2 OS cells). The kinetics of luciferase reporter gene expression after Wnt addition are very similar in an Axin1 and Axin2 knockdown cells. However, if both Axins are knocked down maximal levels of luciferase reporter expression are more than 200 times that of the basal levels.

This unusual behavior of Axin2 is a product of nested feedback circuits as represented in Fig. 4. On the surface it appears that Axin1 is simply redundant, as there is little phenotype from knockdown of either Axin2 if Axin1 is present or vice-versa. But as the detailed dynamical experiments clearly demonstrate Axin2 is a regulator with a unique function. The complete knock down of Axin1 does not achieve the maximal level of β-catenin that can be achieved by inhibiting both Axin1 and Axin2. This gives the impression that Axin2 is more than a backup. Mouse Axin1 and Axin2 show 44% overall identity at the protein level, with higher similarity in domains for ligands and other proteins (11). Yet transgenic replacement in the mouse of the Axin1 coding region for Axin2 or vice versa showed no obvious phenotypes, suggesting that protein sequence differences between Axin1 and Axin2 were of no functional consequence (11). The lack of full redundancy was explained solely by different expression patterns and possibly the lack of Axin2 expression in some cells. The experiments were interpreted as showing that it is the total amount of both Axin proteins that is important for normal function. However, when we looked at these dynamically in a single cell type, differences in both regulation as well differences at the protein sequence level seemed to be important. Such differences may not have been apparent when the assay was normal mouse development, as they may have been suppressed by other mechanisms.

**Fig. 4.**
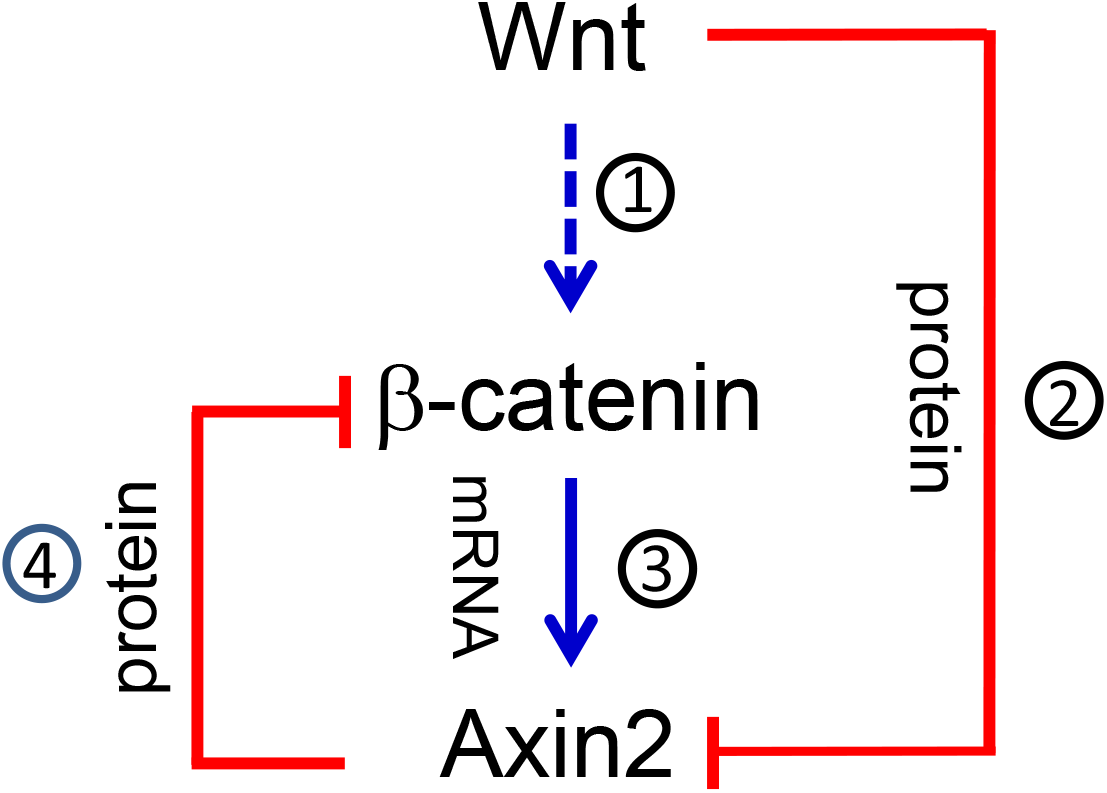
Axin2 and β-catenin feedback regulation in Wnt signaling. (arrow #1) Wnt stimulation stabilizes β-catenin increasing its concentration. (arrow #2) Wnt-3A stimulation destabilizes Axin2 protein. (arrow #3) β-catenin stabilization activates the transcription of Axin2. (arrow #4) Axin2 negatively regulates β-catenin protein expression.

A diagram summarizing the dynamical features of Axin2 is shown in Fig. 4. Wnt increases the levels of β-catenin (Fig. 4, arrow 1) through the inhibition of β-catenin phosphorylation using Axin1 and Axin2 as scaffolds. At the same time, it causes rapid degradation of Axin2 (Fig. 2C, D and E) by some unknown mechanism (Fig. 4, arrow 2). Axin1 behaves differently, decreasing in RKO cells only after 4hrs (Figure 1). The degradation of Axin2 protein is ultimately offset by the increase in Axin2 mRNA and protein driven transcriptionally by the increasing level of β-catenin (Fig. 4, arrow 3) (Fig. 2C and D). The accumulation of Axin2 increases the degradation of β-catenin (Fig. 4, arrow 4). In the case of RKO cells and HEK293T cells the increased degradation of Axin2 is offset by increased transcription and translation of Axin2. For that reason, the knockdown of Axin2 mRNA has little effect on the final level of β-catenin. On the other hand, knockdown of Axin1 is somewhat buffered by Axin2, since a rise in β-catenin will cause higher expression of Axin2, which mitigates the knockdown of Axin1. This might have been more simply engineered by not having Axin2 be both a transcriptional target of β-catenin and a Wnt dependent target of degradation. A clue of a possible biological role for these dynamics is the different effect in U-2OS cells, a human bone derived osteosarcoma line. In this cell line the levels of Axin2 protein overcompensate in response to the Wnt signal (Fig. 2D). There is an initial drop of 60% in Axin2 protein 1 hour after Wnt3A addition but Axin2 protein reaches a 2-fold higher level over the initial level, in response to the much stronger Axin2 transcriptional response, as compared to both the colorectal cell line (RKO) and a cell line derived originally from human embryonic kidney cells (HEK293T) (Fig. 2). The different kinetics of degradation of Axin1 and Axin2 suggest that the sequence differences in these proteins have important biochemical consequences.

There is no way of knowing whether the intricate control of Axin2 is some sort of frozen accident of evolution or some highly selected ingenious mechanisms, but what can be said is that the canonical Wnt pathway is a highly robust largely post-translationally regulated pathway, which operates in many different cell types and in different circumstances. Yet in some circumstances there may be value in being able to adjust the threshold of the response. In the case of bone and tooth development, the Wnt pathway is especially active and here mutations in Axin2 have strong phenotypes; in other circumstances mutations in Axin2 are known to be oncogenic (11). In these cases, regulation can be achieved by simply changing the transcriptional response of Axin2 without changing the core dynamical features of the pathway. The claim of redundancy for the different Axin genes and their intricate regulatory behavior is clearly no longer tenable. It seems more likely that the opportunity to tune the response of the Wnt pathway in different physiological settings provides a reasonable explanation for the distinct regulation of Axin1 and Axin2 and for the feedback circuitry. It may be that many genes, previously deemed genetically redundant, when reevaluated through study of their dynamical features, will reveal important circuitry important in evolution, development, and disease.

## MATERIALS AND METHODS

### Antibodies and reagents

The following antibodies were used: β-catenin (BD Transduction Laboratories), phospho-β-catenin (Ser45) (2564; Cell Signaling), phospho-β-catenin (Ser33/37/Thr41) (Cell Signaling 9561, active β-catenin (anti-ABC) clone 8E7 (Millipore), Axin1 (Cell Signaling; C76H11), Axin2 (Cell Signaling; 76G6), APC (USBiological) and α-tubulin (NeoMarkers).

### Tissue culture and production of Wnt-3A conditioned media

HEK293T (human embryonic kidney), RKO (human colon carcinoma), U-2 OS (human osteosarcoma), L-M(TK-) (mouse fibroblast) and L Wnt-3A [L-M(TK-) cells stably expressing mouse Wnt-3A] were cultured in Dulbecco modified Eagle medium (DMEM) containing 10% fetal calf serum and streptomycin-penicillin in 37°C humidified incubator with 5% CO2. L-Wnt-3A cells were maintained with culture medium supplemented with 0.4 mg/ml G418. All the cell lines were purchased from American Tissue Culture Collection (ATCC), except for RKO cells stably expressing β-catenin activated luciferase reporter, which were kindly provided by Randall T. Moon (University of Washington, Seattle, USA). Wnt-3A conditioned media (Wnt-3A CM) and control media were prepared according to manufacturer’s protocol.

### Treatment with Wnt-3A and MG132

Cultured cells were stimulated with Wnt-3A CM or 100 ng/ml recombinant mouse Wnt-3A (R&D Systems) for the indicated times. MG132 (Calbiochem) treatment was done at a final concentration of 20 µM.

### Preparation of whole-cell protein lysates

The cells were washed twice with ice-cold PBS and lysed with a buffer containing 50 mM Tris/pH 7.6, 150 mM NaCl, 5 mM EDTA/pH 8.0, 0.5% NP-40, supplied with protease inhibitor cocktail (Roche), 1 mM phenylmethylsulfonyl fluoride, and the phosphatase inhibitors sodium fluoride (10 mM), p-nitrophenylphosphate (20 mM), b-glycerophosphate (20 mM), sodium orthovanadate (1 mM), okadaic acid (1 mM), and mycrocystin-LR (1 mM). The lysates were incubated on ice for 0.5 hour, and then cleared by full-speed centrifugation at 4°C for hour. The total protein concentration of the lysates was measured by the Bradford method (BioRad), using bovine serum albumin (BSA) (Sigma-Aldrich) as a standard.

### Qualitative and quantitative immunoblotting

Proteins were resolved by linear SDS-PAGE (7.5% acrylamide). APC was separated by Tris-Acetate 3-8% precast gels (NuPAGE Novex, Invitrogen). The proteins were transferred into nitrocellulose membranes by electrophoretic transfer.

For qualitative immunoblotting, the nitrocellulose membranes were blocked with blocking buffer (TBS, 0.1%Tween-20, 5% nonfat dry milk) for 0.5 hour at room temperature. The membranes were incubated overnight at 4°C with a primary antibody, which was diluted 1:500 to 1:2000. After extensive washing with TBS/0.1% Tween-20, the membranes were incubated for 30 minutes at room temperature with a horseradish peroxidase (HRP)-conjugated secondary antibody (Jackson ImmunoResearch), which was diluted 1: 10,000. The proteins were detected by Enhanced Chemiluminescence (ECL) using ECL Western Blotting Reagents (Amersham, GE Healthcare) or SuperSignal West Dura Extended Duration Substrate (Pierce).

For quantitative immunoblotting, the nitrocellulose membranes were washed with PBS and blocked with Odyssey blocking buffer (Li-Cor). The membranes were incubated overnight at 4°C with a primary antibody, which was diluted 1:500 to 1: 5,000. After extensive washing, the membranes were incubated for 1 hour at room temperature with a fluorescent secondary antibody, which was diluted 1: 10,000. The membranes were then scanned in the appropriate infrared channel using an Odyssey Infrared Scanning System (Li-Cor). The secondary antibodies used were Alexa Fluor 680 donkey anti rabbit (Molecular Probes), and IRDye 800CW goat anti mouse (Li-Cor). The quantification of protein bands was performed using the Odyssey application software (Li-Cor). Protein quantification was calculated on base of measurements of at least three independent experiments in duplicates.

### RNA extraction, cDNA synthesis, and qRT-PCR

RNA was extracted from cells using RNeasy (Qiagen). First strand cDNA was synthesized from 0.5-1 mg RNA by Superscript III First-Strand Synthesis superMix for qRT-PCR (Invitrogen). qRT-PCR reactions were performed using Universal PCR master mix (Applied Biosystems) and TaqMan gene expression assays (Applied Biosystems), according to the manufacturer’s protocol. GAPDH and β-actin were used as endogenous controls, as the expression levels of these genes were found to remain unaffected by Wnt stimulation. Each sample was measured in triplicates. The qRT-PCR reactions were carried out in iCycle iQ (BioRad), as follows: 40 cycles of amplification with 15 second denaturation at 95°C and 1-minute annealing at 60°C. The gene expression analysis was done using iQ5 software (BioRad). Relative and normalized gene expressions were calculated (ΔΔCT). The gene expression was normalized to an endogenous control, such as GAPDH or β-actin, and was relative to a defined control condition.

### β-catenin activated Firefly Luciferase reporter assay

β-catenin activated Luciferase reporter assays in RKO-pBAR cells were performed using a Dual-Luciferase Reporter Assay System (Promega,) and following the protocol provided by the manufacturer. The Firefly and Renilla Luciferase assays included duplicates of each sample and were carried out in white opaque flat bottom 96-well plates (Falcon, Becton Dickinson & Co) using a plate-reader luminometer (Wallac 1420, PerkinElmer). The Luciferase activities were measured for 1 and 3 seconds. The Firefly Luciferase activity was calculated after measuring samples from three independent experiments in duplicates and normalizing the values according to Renilla Luciferase activities in order to correct differences in the amount of cells included in each sample.

## Acknowledgment

We thank Jeremy Gunawardena, Wenzhe Ma, and Bai Luan for critically reading the manuscript and making helpful suggestions. We thank Adriana Rozental for invaluable support, Mike Gage and Leon Peshkin for facilitating completion of the experiments, Randall Moon for providing RKO cells expressing β-catenin activated reporter, Jurgen Behrens for providing Axin2 shRNA plasmid and Yinon Ben-Neriah for Axin1 and APC shRNA plasmids. We thank grants Biochemical studies of mitosis (5R01GM026875-42) and Xenopus Development (5R01HD073104-09) for supporting this work.

## Author contributions

A.R.M. Conceived of the experiments and wrote the paper with the help of M.W.K. A.R.M. performed all of the experiments.

## Data and material availability

All data needed to evaluate the conclusions are present in the paper.

## REFERENCES

1. R. Nusse, Wnt signaling in disease and development. Cell Res. 15, 28–32 (2005).

2. Z. Steinhart, S. Angers, Wnt signaling in development and tissue homeostasis, Development 145 (2018).

3. A. R. Hernandez, A. M. Klein, M. W. Kirschner, Kinetic responses of beta-catenin specify the sites of Wnt control, Science 338, 1337–1340 (2012).

4. M. Gavagan, E. Fagnan, E. B. Speltz, J. G. Zalatan, The scaffold protein Axin promotes signaling specificity within the Wnt pathway by suppressing competing kinase reactions, Cell Syst. 10, 515–525 (2020).

5. E-H Jho, T. Zhang, C. Domon, C-K Joo, J-N Freund, F. Costantini, Wnt/beta-catenin/TCF signaling induces the transcription of Axin2, a negative regulator of the signaling pathway, Mol. Cell Biol. 22, 1172–1183 (2002).

6. J. Y. Leung, F. T. Kolligs, R. Wu, Y. Zhai, R. Kuick, S. Hanash, K. R. Cho, E. R. Fearon, Activation of Axin2 expression by beta-catenin-T-cell factor. J. Biol. Chem. 277, 21657–21665 (2002).

7. B. Lustig, B. Jerchow, M. Sachs, S. Weiler, P. Torsten, U. Karsten, M. van de Wetering, H. Clevers, P. M. Schlag, W. Birchmeier, J. Behrens, Negative feedback loop of Wnt signaling through upregulation of conductin/axin2 in colorectal and liver tumors. Mol Cell Biol. 22, 1184–1193 (2002).

8. H-M I. Yu, B. Jerchow, T-J Sheu, B. Liu, F. Costantini, J. E. Puzas, W. Birchmeier, W. Hsu, The role of Axin2 in calvarial morphogenesis and craniosynostosis. Development 132, 1995–2005 (2005).

9. A. Hlouskova, P. Bielik, O. Bonczek, V. J. Balcar, O. Sery, Mutations in Axin2 as a risk factor for tooth agenesis and cancer: a review. Neuro Endocrinol. Lett. 38, 131–137 (2017).

10. T. L. Biechele, R. T. Moon, Assaying beta-catenin/TCF transcription with beta-catenin/TCF transcription-based reporter constructs. Methods Mol Biol. 468, 99–110 (2008).

11. I. V. Chia, F. Costantini, Mouse axin and axin2/conductin proteins are functionally equivalent in vivo, Mol Cell Biol. 25, 4371–4376 (2005).

12. L. Lammi, S. Arte, M. Somer, H. Jarvinen, P. Lahermo, I. Thesleff, S. Pirinen, P. Nieminen, Mutation in Axin2 cause familial tooth agenesis and predispose to colorectal cáncer, Am J Hum Genet 74, 1043–1050 (2004).

